# CellChat Hotspot: A Targeted Approach to Cell–Cell Communication Analysis With Spatial transcriptomics Validation

**DOI:** 10.64898/2026.11.09.765280

**Authors:** Dario Monaco, Mirea Dioguardi, Maria Dipalma, Eliseo Mattioli, Francesco Alfredo Zito, Francesco Giovannelli, Angela Ricco, Oronzo Brunetti, Antonella Argentiero, Simona De Summa

**Affiliations:** Molecular Diagnostics and Pharmacogenetics Unit, IRCCS Istituto Tumori “Giovanni Paolo II”, Via O. Flacco 65, 70124 Bari, Italy; Pathology Department, IRCCS Istituto Tumori “Giovanni Paolo II”, Via O. Flacco 65, 70124 Bari, Italy; Ospedale San Paolo, Strada Capo Scardicchio, 70132 Bari, Italy; S.S.D. C.O.r.O. Bed Management Presa in Carico, TDM, IRCCS Istituto Tumori “Giovanni Paolo II”, Via O. Flacco 65, 70124 Bari, Italy; Medical Oncology Unit, IRCCS Istituto Tumori “Giovanni Paolo II”, Via O. Flacco 65, 70124 Bari, Italy

**Keywords:** Single-Cell RNA-seq, Cell-Cell Communication, CellChat, Tumor Microenvironment, Spatial Transcriptomics RNA-seq, Colon Cancer

## Abstract

In recent years, single-cell RNA sequencing (scRNA-seq) has enabled the investigation of intercellular communication through Cell–Cell Communication (CCC) tools such as *CellChat*. Despite their potential, these approaches remain challenging to interpret, particularly when global analyses generate complex interaction networks that may include biologically irrelevant signals. To address this limitation, we developed a framework termed *CellChat Hotspot*, designed to improve the biological interpretability of inferred communication networks. This approach integrates gene set enrichment analysis with a context-driven filtering strategy, selecting both genes and cells based on their functional relevance prior to communication inference. Applied to stage II colon cancer data, *CellChat Hotspot* enabled the identification of condition-specific interaction patterns associated with recurrence. Non-recurrent patients showed enrichment of immune-related processes, whereas recurrent cases were characterized by pathways associated with invasion and metastasis. Notably, a loss of interaction in the HSPG pathway between CALB1^+^ and stromal cells was observed in recurrent patients, suggesting a potential disruption of cell–matrix adhesion mechanisms. To verify the reliability of this finding, we applied the framework to spatial transcriptomics data, which confirmed the presence of the *HSPG2–DAG1* interaction in non-recurrent tumors and its near absence in recurrent ones. Overall, this framework provides a biologically guided strategy for studying intercellular communication in complex systems such as the tumor microenvironment.

## 1 Introduction

In recent years, single-cell RNA sequencing (scRNA-seq) has marked a decisive turning point in molecular biology. The ability to analyze gene expression at the level of individual cells has opened new perspectives, enabling exploration of cellular heterogeneity with an unprecedented level of detail [1]. In oncology, in particular, scRNA-seq has enabled a more precise investigation of the cellular dynamics that characterize tumors, contributing to the elucidation of complex mechanisms underlying tumor development [2], relapse risk [3], and treatment response [4]. Alongside the spread of these technologies, there has been rapid development of bioinformatics tools dedicated to their analysis. This constantly evolving landscape requires researchers not only to keep up with methodological advancements but also to develop the ability to address and interpret an increasing level of biological complexity. From the annotation of cellular populations to the identification of robust marker genes, and ultimately to functional characterization, each step of the analysis contributes to building an increasingly accurate representation of the biological system under investigation. In this context, attention has progressively shifted from the study of individual cells to the analysis of interactions between them. This perspective has led to the development of Cell–Cell Communication (CCC) tools [5], a class of methods for inferring cellular communication networks from transcriptomic data. From a methodological standpoint, these approaches mainly rely on two components: on one hand, curated databases containing known ligand–receptor pairs; on the other, mathematical and statistical models used to estimate and prioritize interactions, often through strategies such as cluster label permutation, regularization techniques, and scaling procedures, in order to adapt the inference to the specific characteristics of the analyzed datasets. Cesaro et al [6] reviewed benchmarking analyses focused on CCC tools. CellChat displayed the best performances in terms of usability, robustness, accuracy and, thus, at this stage we focused only on this tool. Despite their great potential, these methodologies still present significant limitations. In particular, the biological interpretation of inferred interactions remains a complex challenge, as it requires deep knowledge of the studied system to distinguish truly functional signals from spurious or poorly informative associations. In light of this complexity, in this work, we focused on the use of CellChat [7], with the aim of improving its biological interpretability and facilitating the identification of the most relevant interactions. To this end, we developed an extended strategy, termed *CellChat Hotspot*, with the aim of better exploiting the biological insights related to recurrence risk of stage II colon cancer patients.

## 2 Data and Methods

### 2.1 Patient cohort and scRNA-seq pre-processing

A cohort of nine stage II colon cancer tissue samples, stratified according to recurrence event, was analyzed. Raw scRNA-seq data were aligned and quantified using Cell Ranger (*v. 7.1.0*). The resulting count matrices were imported into R (*v. 4.5.2*) and converted into Seurat (*v. 5.4*) objects for downstream analysis. Following quality control checks, each dataset was independently normalized using Seurat’s log-normalization method. To correct for batch effects across the nine samples and harmonize expression profiles, we applied Seurat’s anchor-based integration workflow. The resulting integrated Seurat object was used for all downstream analyses. Clustering was performed using the `FindClusters()` function with a resolution of 0.6, resulting in the identification of 17 distinct clusters. These clusters were manually annotated based on marker gene expression identified using the `FindAllMarkers()` function.

### 2.2 Default and Hotspot CellChat analysis

To evaluate communication between cell types, CellChat [7] (*v. 2.2.0*) was used with all cell types and *CellChatDB.human* (*v. 2.0*). CCCs were analyzed through:

1. Default CellChat analysis, following the GitHub guide (CellChat GitHub repository)
2. *CellChat Hotspot*, that was proposed to improve the biological relevance of the results. Thus, three different steps were added before running CellChat:
  1. *The first step was to perform of Gene-set enrichment analysis, using* `AddModule score()` *function, the gene-sets used were Hallmark of cancer [8]. Following the approach described in this work [9], the gene-sets were divided into two categories: Tumor Hallmarks and Tumor Microenvironment Hallmarks. For this analysis, only the Hallmark of the TME (Tumor Microenvironment) was used. Statistically significant differences between conditions were assessed using the Wilcoxon rank-sum test*.
  2. *Cell filtering based on gene set activity: for each gene set, the global median of the enrichment score was computed across all TME cells and used as the activity threshold. For each cell type, the first quantile (Q*_1_*) of the score distribution was then calculated, and a cell type was retained when its Q*_1_ *exceeded the global median, i.e. when at least 75% of its cells scored above the median of the whole TME*.
  3. *Finally, the dataset was restricted to the genes included in the gene set of interest, enabling more targeted analysis of cell communication within the selected biological context*.
  4. *At least CellChat was run with default parameters*.
  5. *CellChat comparative analysis was performed*.

### 2.3 Robustness assessment of CellChat Hotspot

To test whether the communication captured by *CellChat Hotspot* depends on the selected cells a negative control was built by replacing the selected TME populations with the same number of TME cell types showing the lowest *Q* _1_ of the score, excluding previews selected populations. The control was processed with exactly the same pipeline as *CellChat Hotspot*, keeping the four tumor populations and the same gene set.

Significant interactions were extracted with `subsetCommunication` and classified as tumor–tumor, tumor–TME or TME–TME according to the identity of sender and receiver. Because tumor–tumor interactions involve the same cells in both settings, the comparison focused on tumor–TME communication. For each clinical group we compared the number of tumor–TME ligand–receptor interactions, the number of signaling pathways involved and the information flow, defined as the sum of the communication probabilities of the interactions. Information flow was also computed per pathway.

### 2.4 Spatial transcriptomics analysis

Two regions per patient were analyzed with Visium v2 Slide following manufacturer’s protocol and sequenced by NextSeq 550 Illumina platform. blc files were demultiplexed with*bcl2fastq*and the obtained fastq files were analyzed with spaceranger (*v. 3.0.0*, 10X Genomics).

#### Pathologist annotation

Regions of interest (ROIs) were defined by a pathologist, who annotated the Visium spots of each section on the H&E image in Loupe Browser (10x Genomics,*v. 8.0.0*) into histological categories: tumor, stroma, inflammation, smooth muscle and adipose tissue. Spots covering tissue artefacts were annotated as artefact and excluded from all analyses. The annotations were exported as per-spot barcode tables and matched to the Visium data by spot barcode.

#### Spot deconvolution

The cell-type composition of each Visium spot was estimated with CARD [10], (R package *v. 1.1*). The reference was the scRNA-seq dataset described above. Raw UMI counts from the two count layers of the reference Seurat object (stage II with and without recurrence) were merged on their shared genes, and each patient was declared as a separate sample (`sample.varname`), so that CARD builds the cell-type reference basis while accounting for variability between patients. All annotated cell types were used (`ct.select`).

The spatial input of each section was the raw count matrix of the Spatial assay and the spot coordinates (Seurat `GetTissueCoordinates`). Each Visium v2 section was deconvolved independently. CARD was run with its default quality-control settings (*minCountGene*=100, *minCountSpot*=5), and with its default selection of informative cell-type marker genes and default spatial model parameters.

#### Spot assignment

The CARD proportions of each spot were first row-normalised to sum to 1(missing or negative values set to 0). A spot was assigned its most abundant cell type only when that proportion was at least 0.10 and exceeded the second most abundant by at least 0.05. Spots failing either criterion were labelled mixed (unassigned), and spots removed by CARD quality control were labelled not deconvolved; both were excluded from label-based analyses. Within a section, cell types with fewer than 10 assigned spots were not tested in neighbourhood enrichment and were not used as communication groups in CellChat.

#### CellChat

Each spot was labelled with its dominant cell type from the full-annotation CARD deconvolution, using the rule described above (row-normalised proportion ≥ 0.10 and ≥ 0.05 above the second most abundant type). Unassigned spots were removed. Each spot also received its ROI label from the pathologist’s annotation; spots left unannotated were assigned a section-specific default region and spots annotated as artefact were excluded. Communication groups were defined as cell type × ROI combinations (e.g. mCAF in stroma), so that a single model captured both communication within a region and communication across region borders. Groups with fewer than 10 spots were discarded.

Ligand–receptor communication was inferred with CellChat [7], (R package *v. 2.2.0*). Each Visium section was analysed independently, with a single CellChat model per whole section. Raw counts of the Spatial assay and coordinates were used as input, with the spatial scale factors set as recommended by CellChat for Visium (spot size 65 *µm*, converted to pixels with the section’s `spot_diameter_fullres`, and a tolerance of half the spot size). The complete human ligand–receptor database (*CellChatDB.human*) was used. Over-expressed genes and interactions were identified with default settings. Communication probabilities were computed with the truncated mean (*type = “triMean”*), with distance constraints (*distance.use = TRUE, interaction.range =*250*µm, scale.distance*=0.01) and with contact-dependent signalling restricted to spots within 100*µm*(*contact.dependent = TRUE, contact.range = 100*). Interactions involving groups with fewer than 10 spots were filtered out (*filterCommunication, min.cells = 10*). Ligand–receptor pairs with permutation *p* < 0.05 and communication probability > 0 were considered significant. Communication was then summarised at the pathway level (`computeCommunProbPathway`), aggregated into networks (`aggregateNet`), and sender and receiver roles were computed with network centrality analysis (`netAnalysis_computeCentrality`).

Each significant interaction was classified as within-ROI when the sender and receiver groups belonged to the same region, and as cross-ROI otherwise. Pathway strength, the number of significant interactions and the outgoing and incoming communication of each cell type were summarised for each ROI and for each ordered pair of ROIs.

#### Spatial validation of the HSPG2–DAG1 interaction

The interaction prioritised by *CellChat Hotspot* in the scRNA-seq data, HSPG2–DAG1 (HSPG pathway, ECM– receptor), was tested in the Visium sections of the same nine patients:15 sections in total,7 from the five patients without recurrence and 8 from the four patients with recurrence.

For each section, all sender–receiver edges of HSPG2–DAG1 with communication probability > 0 were extracted from the whole-section CellChat model described above, pooling the region of the sender and receiver groups. For every pair of sender and receiver cell types, each section was classified as significant (*p* <0.05), detected but not significant, tested with probability 0(both groups present), or not testable (at least one cell type had fewer than 10 assigned spots and was not a CellChat group). For the condition-level network, the probabilities of significant edges were summed for each pair of cell types across regions and averaged over the sections of each condition (sections without the edge counted as 0).

##### Ligand–receptor proxy

To quantify the interaction independently of CellChat and of the dominant-type assignment, a spot-level ligand–receptor proxy was computed from the CARD proportions. Raw counts of the Spatial assay were normalised to

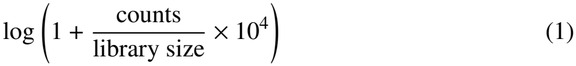

For spot i, with *HSPG2* expression *L*_*i*_, *DAG1* expression *R*_*i*_ and CARD proportions *p*:

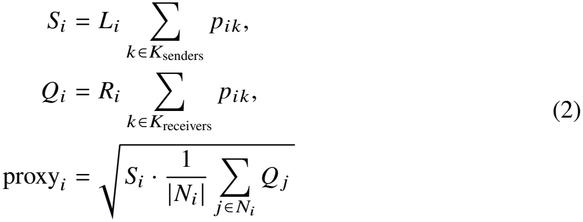

Where *N*_*i*_ is the set of spots (including i) whose centre lies within 250 *µm* of spot i, the interaction range used in CellChat for this ECM–receptor pair. Distances were computed from the full-resolution pixel coordinates with the same scale as CellChat (65 *µm* per spot diameter). The proxy is greater than zero only where a spot with ligand-expressing senders has *DAG1-expressing* receivers nearby. Only deconvolved spots in the tumor and stroma regions were used for the comparison. The patient was the unit of analysis. For each patient (sections pooled) we computed the percentage of spots with proxy > 0, the mean proxy over all spots and the median proxy of positive spots. Patients without and with recurrence (5 vs 4) were compared. As a complementary spot-level analysis, a linear mixed model with patient and section as nested random intercepts, `proxy ~ recurrence + (1 | patient/section)`, was fitted with lme4 and tested with lmerTest.

## 3 Results

### 3.1 scRNA dataset

The scRNA-seq dataset used in this study consists of nine stage II colon cancer patients, including five patients without recurrence and four patients who experienced disease recurrence. The UMAP representation (Figure 1A) illustrates the overall cellular landscape, highlighting both the distribution of the two clinical conditions and the identification of 17 distinct cellular populations. The relative proportions of the identified cell populations (Figure 1B) reveal a high degree of inter-condition heterogeneity. This variability underscores the complex cellular composition of the tumor microenvironment and supports the need for more refined analytical approaches to dissect condition-specific biological signals. This complexity motivated the application of CCC analyses to uncover condition-specific interaction patterns.

**Fig. 1.**
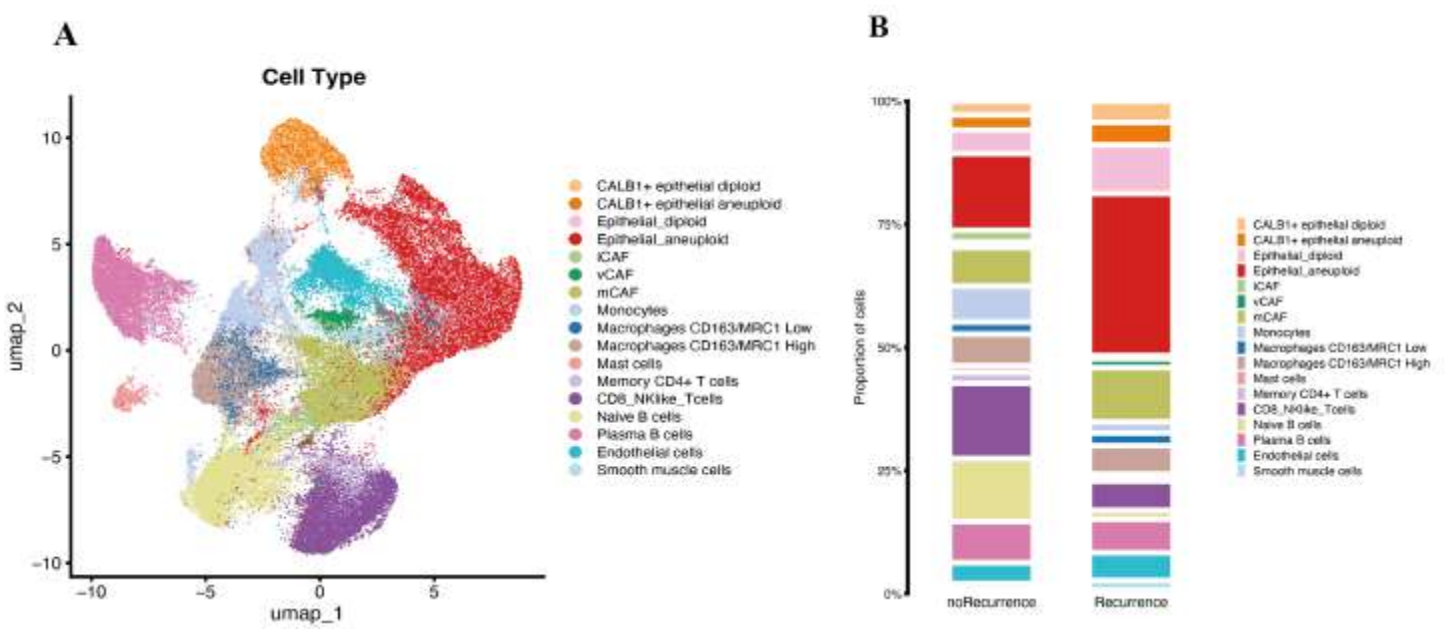
Distribution of single-cell dataset: A. UMAP distribution of the 17 cell types split by Recurrence status; B. Stacked barplot that illustrates the proportions of cell types.

### 3.2 Global CellChat

To characterize the overall intercellular communication landscape, we applied CellChat to the integrated single-cell dataset. The analysis revealed a complex, highly interconnected network of signaling interactions across the 17 identified cellular populations (Figure 2A).

**Fig. 2.**
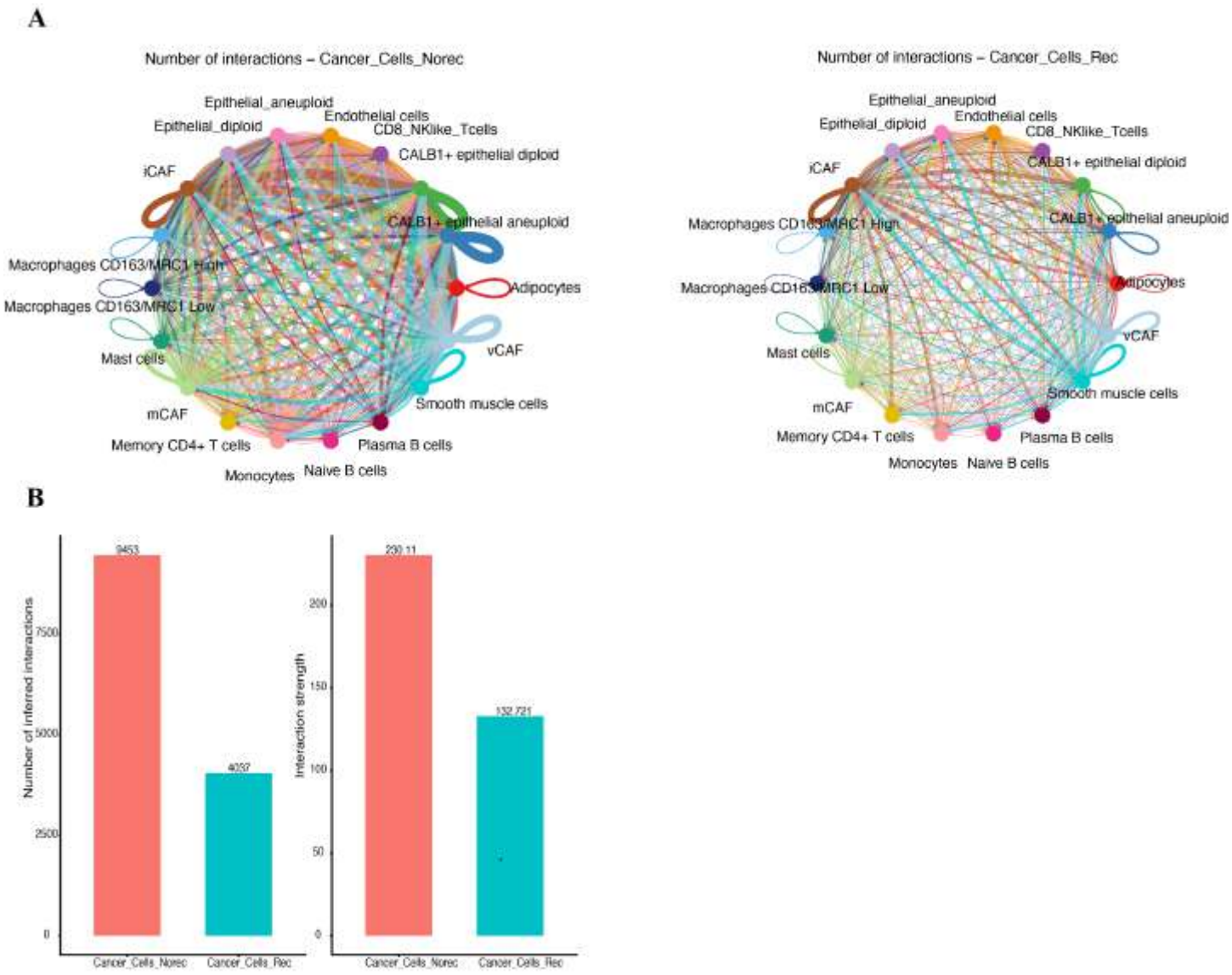
Global CellChat: A. Circle plots that show the interactions between the different populations; B. Difference between the two conditions in terms of number of interactions and strength.

At the global level, it is possible to appreciate an increase in the number of communications and in the strength of interactions among non-recurrence patients (Figure 2B).

However, despite these differences, the global CellChat analysis remains limited in its ability to resolve fine-grained, functionally relevant interactions.

To overcome this limitation, we next applied the *CellChat Hotspot* approach to focus on biologically relevant and context-specific signaling interactions.

### 3.3 CellChat Hotspot

The gene set activity analysis revealed distinct enrichment patterns between the two clinical conditions. Specifically, gene sets associated with activation of invasion and metastasis were enriched in patients with recurrence, whereas non-recurrent patients showed higher enrichment for avoiding immune destruction and tumor-promoting inflammation (Figure 3A).

**Fig. 3.**
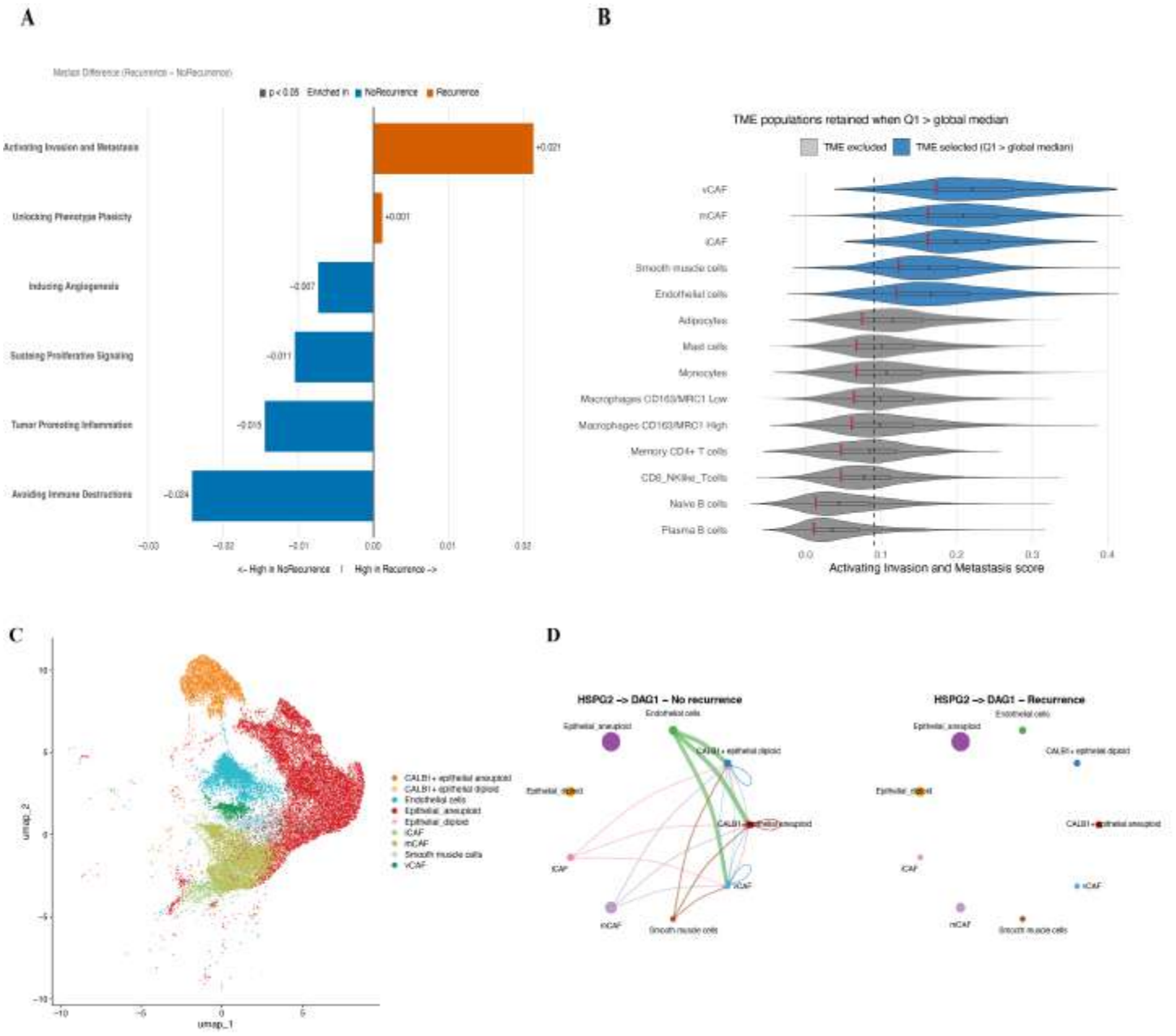
CellChat Hotspot: A. Comparison of Hallmark AddModuleScore() results; statistical analysis was performed using the Wilcoxon test; B. Violin plot showing the cells of the TME responsible for this enrichment; C. UMAP showing the cells responsible for the gene set enrichment; D. Differential interaction of HSPG2 and DAG1 (respectively, ligand and receptor) pathway between cells.

The cutoff applied to the Activating invasion and metastasis gene set highlighted tumor-related and stromal populations, including epithelial cells (both CALB1^+^ and non-CALB1^+^), endothelial cells, smooth muscle cells, and cancer-associated fibroblasts (Figure 3B-C).

CellChat was the first algorithm to be able to perform comparative analysis between conditions contrasting interaction numbers, interaction strengths, signaling information flow, sender/receiver roles, and network topology across conditions. Comparative CellChat analysis identified an *HSPG2-DAG1* (pathway HSPG) interaction between CALB1^+^ epithelial cells and stromal cells (mCAF, vCAF, iCAF, endothelial cells and smooth muscle cells) in the no-recurrence condition, whereas this interaction was not retained in the recurrence communication network (Figure 3D).

### 3.4 Robustness assessment of CellChat Hotspot

We then assessed the robustness of the method on the same dataset, starting from the number of inferred interactions. All interactions identified by *CellChat Hotspot* were also present in the default CellChat output (Figure 4A), indicating that the method does not introduce new interactions. Compared with default CellChat, Hotspot reduced the number of interactions by approximately 78% in non-recurrent patients (2,095 of 9,453 retained) and by approximately 76% in recurrent patients (960 of 4,037 retained).

**Fig. 4.**
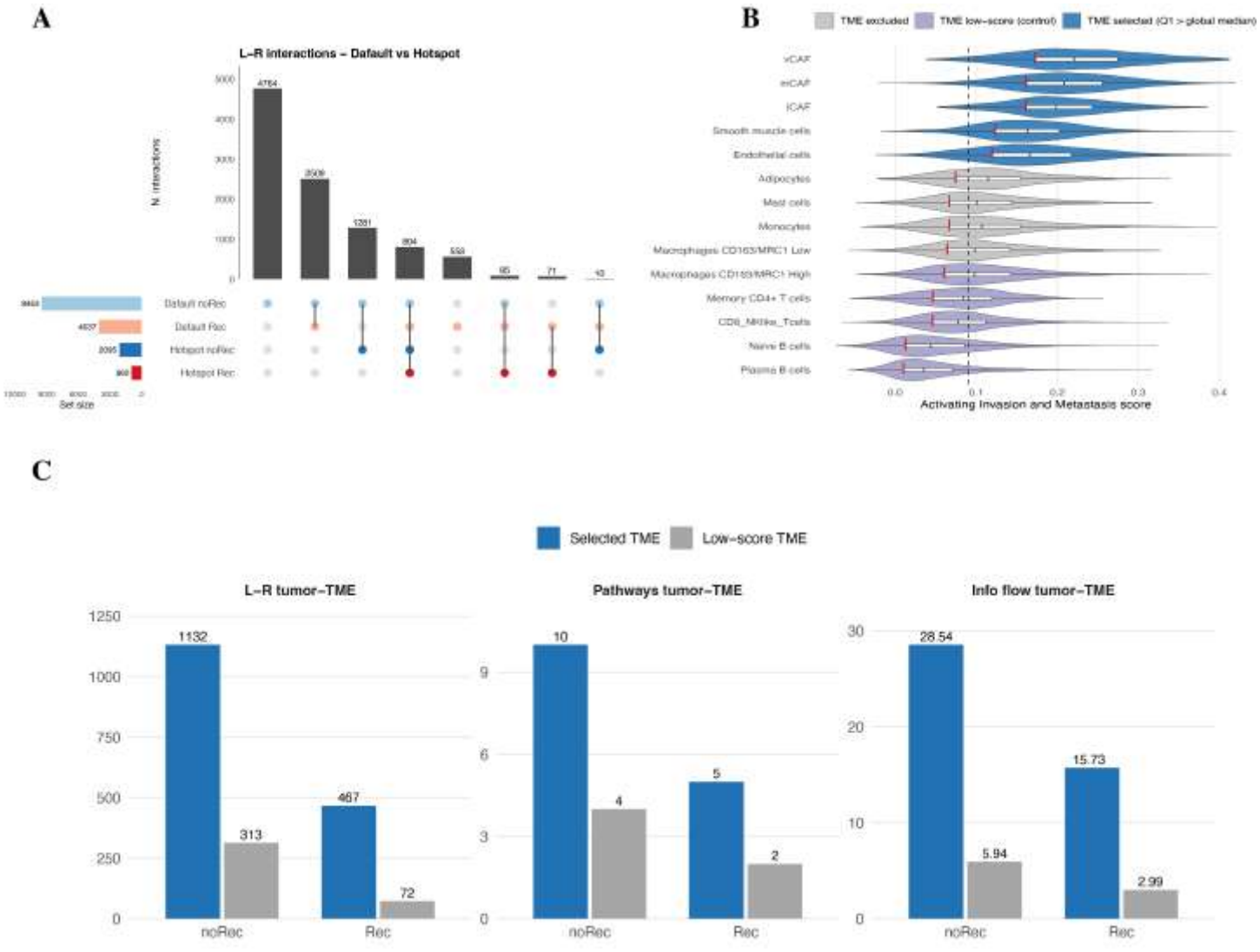
Robustness assessment of CellChat Hotspot: A. UpSet plot of ligand–receptor interactions inferred by default CellChat and CellChat Hotspot in non-recurrent and recurrent patients; B. Gene-set score across TME cell types: selected (blue) and low-score control (purple) populations; dashed line, global median; C. Tumor–TME interactions, pathways and information flow obtained with selected versus low-score TME populations.

To test whether the inferred communication depended on cell selection rather than on gene restriction alone, we repeated the analysis replacing the selected TME populations with the same number of TME populations with the lowest scores, while keeping tumor populations and the gene set unchanged (Figure 4B). Low-score populations yielded 3.6-fold (non-recurrent) and 6.5-fold (recurrent) fewer tumor–TME interactions and an approximately 5-fold lower information flow, showing that the signals captured by *CellChat Hotspot* depend on score-based cell selection (Figure 4C).

### 3.5 Spatial validation of HSPG2–DAG1 signaling

To validate the stromal–tumor HSPG2–DAG1 interaction identified by *CellChat Hotspot*, we analyzed spatial transcriptomics data from the whole cohort, including five non-recurrent and four recurrent patients. Spot-level cell-type composition was inferred by deconvolution using the scRNA-seq annotation as reference. As described in Methods section, CellChat was run on the whole section taking into account pathologist’s annotation to be sure to identify stroma-tumor CC interactions and not only intra-region interactions. Figure 5A–B shows one representative non-recurrent and one recurrent patient, in which CALB1^+^ epithelial cells mapped to tumor regions and CAFs and endothelial cells to the surrounding stroma.

**Fig. 5.**
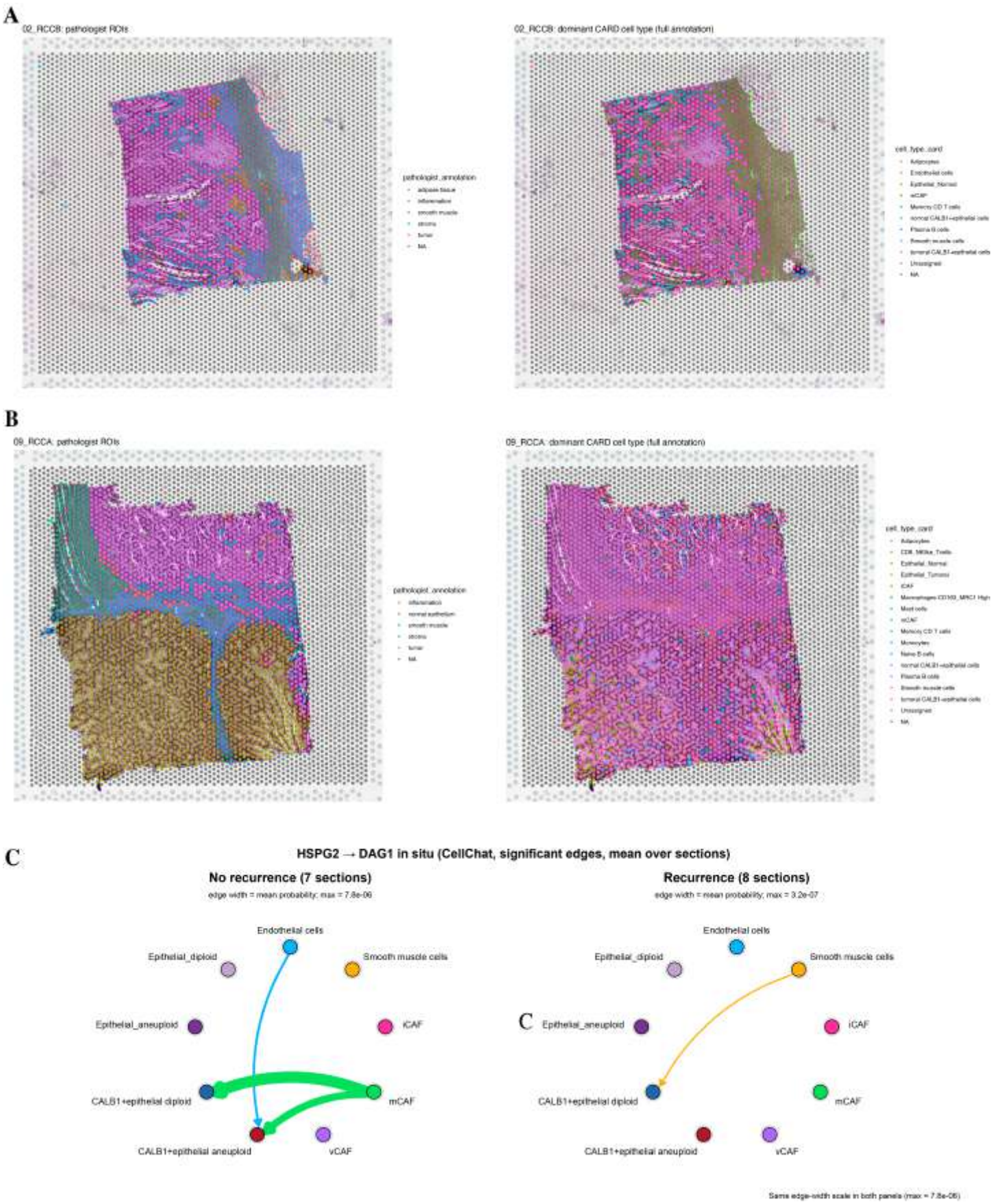
Spatial cell-type deconvolution and in situ HSPG2→DAG1 communication: (A–B) Representative ROIs from a non-recurrent and a recurrent patient: dominant cell type per spot inferred by CARD deconvolution (left) and pathologist annotation of tissue regions (right); C. Circle plots of significant HSPG2→DAG1 interactions inferred by spatial CellChat analysis in non-recurrent and recurrent patients.

Spatial CellChat analysis across the cohort confirmed the pattern observed in the scRNA-seq data (Figure 5C). In non-recurrent tumors, significant HSPG2→DAG1 interactions were detected from mCAFs to CALB1^+^ epithelial cells and from endothelial cells to CALB1^+^ aneuploid epithelial cells. In recurrent tumors, the interaction was almost completely lost, with only a weak signal retained from smooth muscle cells to CALB1^+^ diploid cells. Compared with the scRNA-seq analysis, in which endothelial cells were the main source of *HSPG2*, mCAFs emerged as the dominant sender in situ, likely reflecting their abundance in the stroma adjacent to tumor regions.

To visualize the interaction in tissue context, we mapped *HSPG2* and *DAG1* expression, a spot-level ligand–receptor proxy (250 *µm* neighborhood) and the significant CellChat edges onto the histological sections of the representative patients. In the non-recurrent patient, *HSPG2* and *DAG1* were co-expressed in a large fraction of spots across tumor and stromal regions, and four significant HSPG2→DAG1 edges were detected in both sections, all originating from mCAFs located either in the stroma or within tumor regions and directed to CALB1^+^ diploid and aneuploid epithelial cells (Figure 6A). In contrast, in the recurrent patient co-expression was sparse and no significant edge was detected in either section (Figure 6B).

**Fig. 6.**
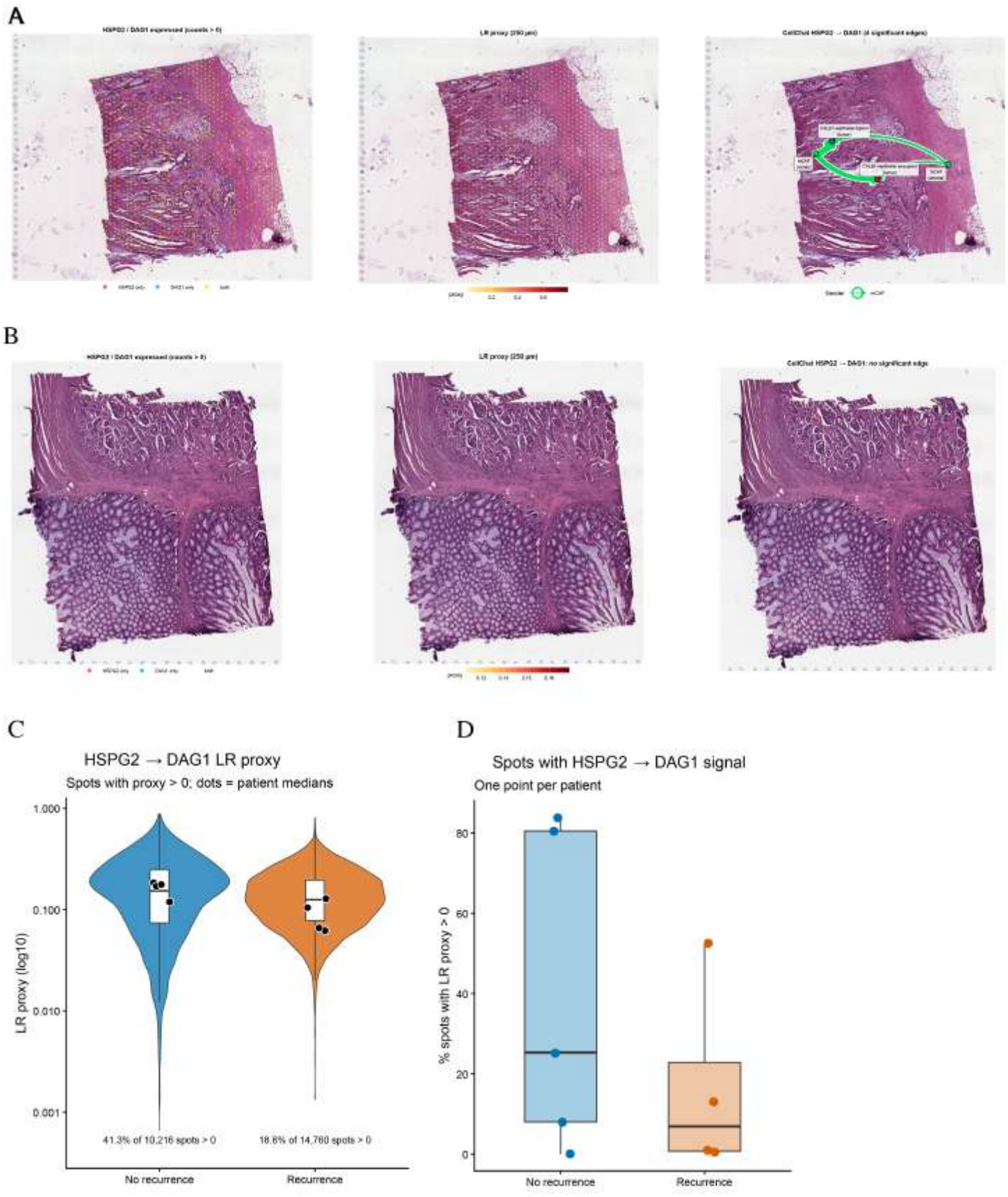
In situ mapping and quantification of HSPG2→DAG1 signaling: Representative ROIs from a non-recurrent (A) and a recurrent (B) patient: spots expressing *HSPG2, DAG1* or both (left), spot-level ligand–receptor proxy (middle), and significant CellChat HSPG2→DAG1 edges (right); C. Distribution of the HSPG2→DAG1 proxy in signal-positive spots in non-recurrent and recurrent samples; D. Percentage of spots with proxy > 0 per patient.

To further confirm the HSPG2→DAG1 interaction, we calculated LR-proxy which is independent both from CellChat analysis and cell type assignment. In tumor and stroma regions, 41.3% of 10,261 spots from patients with recurrence (Figure 6C). Among positive spots, the proxy had a similar range in the two groups, and patient medians largely overlapped (Figure 6C). The difference therefore lay mainly in how much of the tissue carried the signal rather than in its intensity. At the patient level the percentage of positive spots was higher without recurrence (median 71% vs 25%, Figure 6D).

Overall, the spatial data support the single-cell finding: HSPG2–DAG1 communication from stromal cells to CALB1^+^ epithelial cells is present in situ and is more extensive in tumors that did not recur, although the small number of patients and the heterogeneity between them do not allow a formal test of the association with recurrence.

## 4 Discussion

This study highlights how the analysis of cell–cell communication remains inherently complex, requiring a high level of biological knowledge and careful interpretation. This complexity becomes particularly evident when investigating the tumor microenvironment, especially in the absence of clearly defined a priori hypotheses or specific cellular populations of interest. In such contexts, global approaches may generate extensive interaction networks that are difficult to interpret, as observed in our global CellChat analysis. To address this limitation, we developed an extended strategy, termed *CellChat Hotspot*, designed to improve the biological interpretability of inferred communication networks. By integrating gene set enrichment analysis, cell filtering based on functional activity, and targeted gene selection prior to communication inference, this approach enables a more focused identification of relevant interactions and reduces the impact of spurious or less informative signals.

Applied to stage II colon cancer samples, the *CellChat Hotspot* framework enabled the identification of condition-specific signaling patterns associated with recurrence and non-recurrence states, highlighting differences that were not clearly resolved by a global analysis alone. In particular, gene set enrichment analysis revealed distinct functional behaviors between the two conditions, with non-recurrent patients showing enrichment of immune-related processes, consistent with a more active immune-associated microenvironment, whereas recurrent patients displayed enrichment of pathways related to invasion and metastasis.

These findings were further supported by the *CellChat Hotspot* analysis, which identified specific alterations in intercellular communication. Notably, HSPG2–DAG1 signaling from stromal populations to CALB1^+^ epithelial cells was lost in recurrent patients. Perlecan (HSPG2), a major membrane basement proteoglycan [11], binds *α*-dystroglycan (DAG1) [12] and anchors epithelial cells to the basement membrane, contributing to the maintenance of epithelial polarity. Loss of this cell–matrix interaction is consistent with detachment from the basement membrane [13] and with an epithelial–mesenchymal transition (EMT)-like state, which is linked to tumor progression and recurrence in colorectal cancer [14].

Spatial transcriptomics provided an independent validation of these findings in tissue, confirming the presence of HSPG2–DAG1 interactions in non-recurrent tumors and their reduced extent in recurrent ones. However, some limitations of the spatial analysis should be considered. Each spot captures the transcriptome of multiple neighboring cells, so interactions cannot be resolved at single-cell level. In addition, each spot was assigned to its most abundant cell type after deconvolution, which favors large stromal populations over less abundant ones. This likely explains why mCAFs, rather than endothelial cells as in the scRNA-seq data, emerged as the dominant HSPG2 sender in situ. This limitation was partly addressed by the LR-proxy, which uses the full CARD proportions of each spot rather than its dominant cell type, and is therefore independent of both the cell-type assignment and the CellChat analysis.

Overall, this approach provides a flexible and biologically guided framework for studying intercellular communication, particularly in complex systems such as the tumor microenvironment, and may support more robust hypothesis generation and patient stratification in future studies. Moreover, since cell and gene selection are performed before communication inference, the framework could be readily applied to other CCC tools based on ligand–receptor databases.

## Acknowledgments

We thank all the patients who participated in this study. The single-cell RNA-seq and spatial transcriptomics data were generated with funding from the Italian Ministry of Health (Ricerca Corrente 2025).

